# Active information sampling varies across the cardiac cycle

**DOI:** 10.1101/283838

**Authors:** Stella Kunzendorf, Felix Klotzsche, Mert Akbal, Arno Villringer, Sven Ohl, Michael Gaebler

## Abstract

Perception and cognition oscillate with fluctuating bodily states. For example, visual processing has been shown to change with alternating cardiac phases. Here, we study the heartbeat’s role for active information sampling—testing whether humans implicitly act upon their environment so that relevant signals appear during preferred cardiac phases.

During the encoding period of a visual memory experiment, participants clicked through a set of emotional pictures to memorize them for a later recognition test. By self-paced key press, they actively prompted the onset of shortly (100-ms) presented pictures. Simultaneously recorded electrocardiograms allowed us to analyse the self-initiated picture onsets relative to the heartbeat. We find that self-initiated picture onsets vary across the cardiac cycle, showing an increase during cardiac systole, while memory performance was not affected by the heartbeat. We conclude that active information sampling integrates heart-related signals, thereby extending previous findings on the association between body-brain interactions and behaviour.

## Introduction

We perceive and act upon the world while our brain continuously integrates exteroceptive and interoceptive information, that is, information received from external (e.g., through vision and touch) and internal sources (e.g., through viscerosensation and proprioception), respectively (Barrett & Simmons, 2015; Kleckner et al., 2017). Through the fine-tuned interplay of brain and body, we are able to react to changes in the external and the internal environment to maintain or restore our bodily integrity and well-being. Via feedback loops, the brain thereby receives afferent information about bodily states to regulate and adjust bodily activity accordingly (Craig, 2002; Critchley & Harrison, 2013; Mayer, 2011; Saper, 2002). To fully capture the bi-directionality of brain-body interactions and their association with mental processes, it is essential to investigate if and how such afferent bodily information modulates our thoughts, feelings, and behaviour.

One approach to study bodily influences on cognition and behaviour exploits natural physiological fluctuations. Such fluctuations occur at multiple time scales—ranging from milliseconds to weeks (e.g., brain oscillations, heartbeats, the circadian rhythm, or the menstrual cycle)—and they dynamically interact with each other as well as with the environment (Glass, 2001). How such natural physiological variability is processed in the brain remains poorly understood. However, it has been shown that brain and beyond-brain organ systems (e.g., the cardiorespiratory or the gastrointestinal system) co-vary in their oscillatory activity (Fan et al., 2012; Luft & Bhattacharya, 2015; Richter, Babo-Rebelo, Schwartz, & Tallon-Baudry, 2017; Thayer, Åhs, Fredrikson, Sollers, & Wager, 2012).

Particularly the heart, as a fundamental internal oscillator, has been the target of a growing body of research that investigates how cardiac fluctuations are integrated with the processing of external stimuli (Critchley & Garfinkel, 2018). Cardiac activity occurs in a cycle of two phases: During diastole, the ventricles relax to be filled with blood; during systole, the ventricles contract and eject blood into the arteries, while visceral pathways send information about each heartbeat to the brain (Critchley & Harrison, 2013). Such natural phasic changes of the cardiovascular state have been mainly associated with variations in perception: For sensory processing, which is typically measured with detection tasks or reaction time tasks, response to passively presented stimuli has been shown to be attenuated during early cardiac phases (i.e., during systole) or relatively enhanced at later time points in the cardiac cycle (i.e., at diastole) (Birren, Cardon, & Phillips, 1963; Callaway & Layne, 1964; Edwards, Ring, McIntyre, Carroll, & Martin, 2007; Lacey & Lacey, 1974; McIntyre, Ring, Edwards, & Carroll, 2008; Réquin & Brouchon, 1964; Saari & Pappas, 1976; Sandman, McCanne, Kaiser, & Diamond, 1977; Wilkinson, McIntyre, & Edwards, 2013).

A growing number of more recent findings, however, suggests *facilitated* processing during systole, specifically for task-or context-relevant stimuli. While enhanced processing during systole was also reported for non-emotional visual stimuli (Pramme, Larra, Schächinger, & Frings, 2014, 2016), an emotional specificity of this effect was observed when testing neutral stimuli vs. valenced stimuli like emotional faces—particularly for emotionally arousing fear or threat stimuli (Azevedo, Badoud, & Tsakiris, 2018; Azevedo, Garfinkel, Critchley, & Tsakiris, 2017; Garfinkel et al., 2014). Thus pointing towards an increase in emotional salience through interoceptive channels (Critchley & Harrison, 2013), these findings correspond with evidence for preferential stimulus processing (e.g., enhanced perception and memory) fostered by states of general psychophysiological arousal (Cahill & McGaugh, 1998; Mather, Clewett, Sakaki, & Harley, 2016; Mather & Sutherland, 2011; McGaugh, 2015; Tambini, Rimmele, Phelps, & Davachi, 2017). At the same time, phasic cardiac modulation of stimulus processing has been associated with altered memory formation and retrieval (Fiacconi, Peter, Owais, & Köhler, 2016; Garfinkel et al., 2013)—also in the context of respiratory oscillations (Zelano et al., 2016). Taken together, sensory processing is differentially modulated during early cardiac phases, indicating a selective processing benefit for relevant (e.g., emotionally arousing) stimuli, while other perceptual processes are attenuated (Garfinkel & Critchley, 2016). This suggests that cardiac (or cardio-respiratory) fluctuations not only are an important target of efferent arousal regulation but contribute to afferent signalling of bodily arousal states to the brain (Critchley & Harrison, 2013).

However, these studies investigating cardiac influences on perception and cognition have only employed passive stimulus presentation, which ignores *self-initiated action* as a crucial dimension of sensory and particularly visual processing. Mediating our engagement with a visual scene, motor actions dynamically orchestrate incoming sensory data and thus strongly influence visual perception—selecting what information is preferentially processed (Benedetto, Spinelli, & Morrone, 2016; Tomassini, Spinelli, Jacono, Sandini, & Morrone, 2015). Sensorimotor coupling has also been linked to periodic attentional fluctuations (Hogendoorn, 2016; Morillon, Schroeder, & Wyart, 2014). For example in the visual domain, saccadic eye movements are preceded by a shift of attention to the saccade target resulting in strongly improved visual performance (Deubel & Schneider, 1996; Kowler, Anderson, Dosher, & Blaser, 1995; Li, Barbot, & Carrasco, 2016; Ohl, Kuper, & Rolfs, 2017) and memory performance at the saccade target location (Hanning, Jonikaitis, Deubel, & Szinte, 2016; Ohl & Rolfs, 2017) with corresponding neural enhancement in early visual cortex (Merrikhi et al., 2017; Moore, Tolias, & Schiller, 1998). There is sparse evidence that connects the heartbeat to general action generation and the few studies that investigated if movements are modulated across the cardiac cycle indicate that systole provides a facilitating time window for spontaneous motor activity—both in the somatomotor (Mets, Konttinen, & Lyytinen, 2007) as well as in the oculomotor domain (Ohl, Wohltat, Kliegl, Pollatos, & Engbert, 2016).

Based on findings of facilitated processing for visual stimuli (Azevedo et al., 2018, 2017; Garfinkel et al., 2014; Pramme et al., 2014, 2016) and increased oculomotor activity (Ohl et al., 2016) during early phases of the cardiac cycle, we here hypothesized that active information sampling (i.e., self-initiated action towards a visual stimulus) shows periodic variations with the phase of our heartbeat. To investigate perception and action within a comprehensive framework of mind-brain-body interactions, we here studied cardiac-related sensorimotor processing in a self-paced visual sampling paradigm, in which participants decide when to press a key to see a task-relevant visual stimulus. Extending studies that emphasize a selectivity of this effect for motivationally salient, passively presented stimuli (Azevedo et al., 2018, 2017; Garfinkel et al., 2014), we predicted that observers implicitly act upon a relevant visual stimulus such that it is received (and perceived) during preferred cardiac phases. More specifically, we hypothesized that visual sampling would be biased towards processing task-relevant pictures during systole.

We assessed the emotional specificity of cardiac-phase effects (Garfinkel et al., 2014; Garfinkel & Critchley, 2016) by presenting negative, positive, and neutral pictures (cf. **Methods**). To further induce stimulus relevance, participants were instructed to memorize the pictures during sampling and their recognition memory was tested after a delay of several minutes. This enabled us to also address the previously reported link between memory performance and the cardiac cycle: Based on the abovementioned systolic modulation of memory formation (Garfinkel et al., 2013), we expected recognition performance to be modulated by the cardiac timing of memory probes during encoding. Specifically, we hypothesized that memory performance for pictures encoded at different time points of the cardiac cycle is not equally distributed, but varies across the cardiac cycle.

## Methods

### Preregistration

The protocol and the hypotheses of our study were pre-registered prior to the data acquisition using the Open Science Framework (https://osf.io/5z8rx/).

### Participants

47 (23 female) healthy, young, right-handed subjects (age: 18 – 34 years, M = 25.8 years, SD = 4.31) with normal or corrected-to-normal vision participated in this study. Four subjects were excluded due to deviant cardiovascular parameters: two subjects with tachycardic mean resting heart rates (> 100 bpm), parallel to previous studies (Edwards, McIntyre, Carroll, Ring, & Martin, 2002; Garfinkel et al., 2014; Wilkinson et al., 2013); one subject with hypertonic blood pressure (171/89 mmHg), based on Tukey’s (1977) criterion of 1.5 times the interquartile range (IQR) above the third quartile (Q3 = 122 mmHg, IQR = 20.5 mmHg); one subject with numerous ventricular extrasystoles during the experimental period (> 10 per minute). The sample size was based on previous cardiac cycle studies (mainly Fiacconi et al., 2016): We aimed for a net sample size of 40 to enter the analyses (cf. https://osf.io/5z8rx/) expecting 10% participant exclusions. Participants were recruited through the ORSEE-based (Greiner, 2015) participant database of the Berlin School of Mind and Brain and received a monetary compensation of 9 €/h for their participation. All participants were naïve regarding the purpose of the study and signed informed consent before participation. The study followed the Declaration of Helsinki and was approved by the Ethics Committee of the Department of Psychology at the Humboldt-Universität zu Berlin.

### Setup and experimental task

(cf. **Fig. 1a**) Participants were seated in a dimly lit room in front of a gamma-linearized 19-inch Cathode ray tube (CRT) monitor (Samsung Syncmaster 959NF, Suwong, Korea) with a refresh rate of 100 Hz and a spatial resolution of 1280×1024 pixels. Their head was positioned on a chin rest at a distance of 50 cm from the screen. The participants’ task comprised two parts: During the *encoding* period, participants were asked to click through a picture set (800×600 pixels) in self-paced speed and to memorize the pictures for a subsequent memory test. By button press, they prompted the immediate onset of the next picture, which appeared for 100 ms. In between self-chosen key presses (i.e., picture onsets), a central fixation cross was presented. After a break of five minutes, they completed the *recognition* period, during which they indicated for each picture whether or not they had seen it before. Here, pictures were passively presented for 100 ms, followed by a centrally presented fixation cross until participants entered their recognition response (“old”, “new”) via key press.

**Figure 1.**
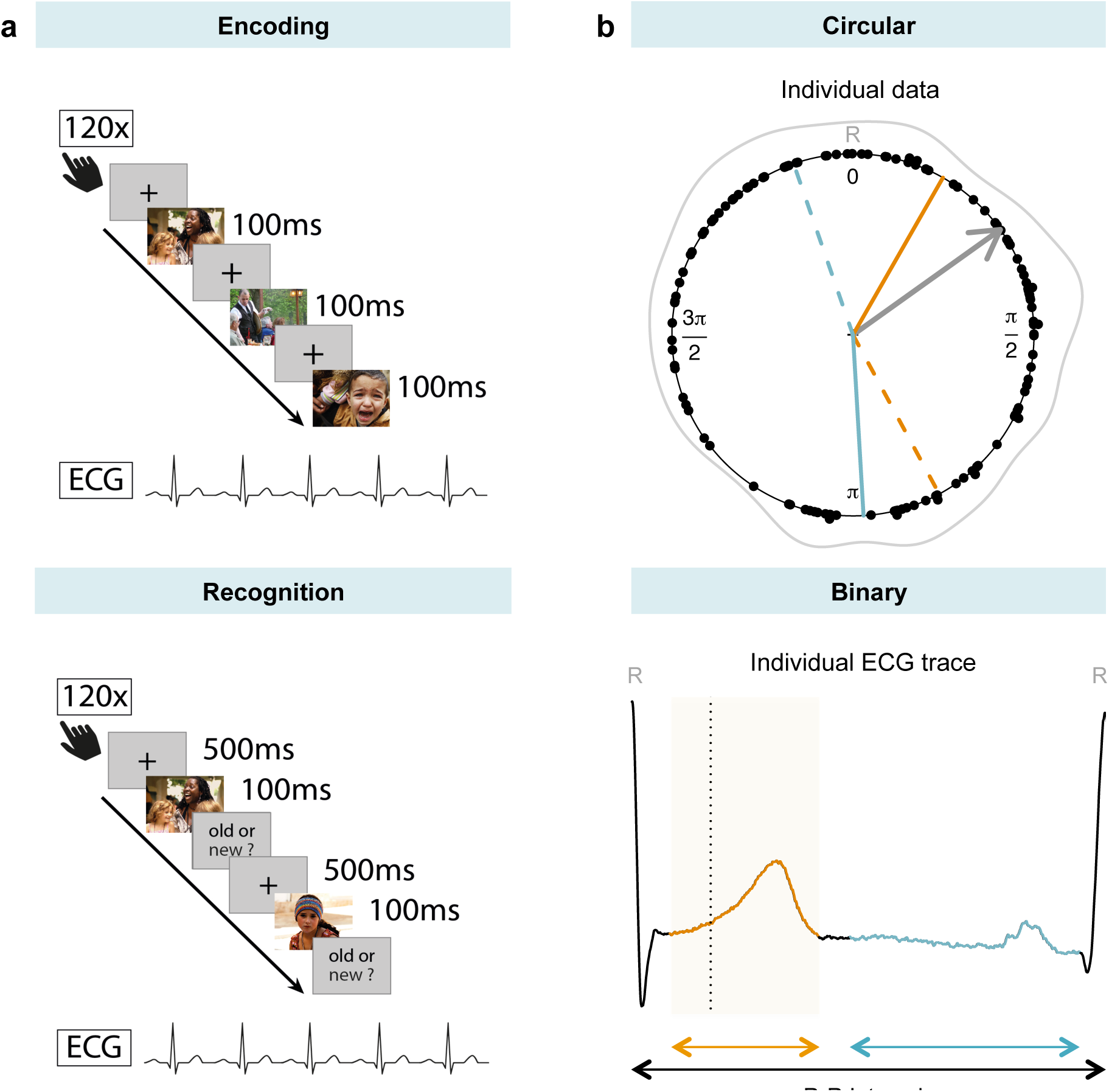
Experimental setup and data analysis. ***a***, During encoding (top), participants prompted by button press the onset of the next picture, which could be positive, neutral, or negative. In the recognition period (bottom), they indicated for each picture (60 old, 60 new) whether or not they had seen it before. Simultaneous ECG was recorded to analyse behaviour relative to the cardiac cycle: ***b***, For circular analysis (top), values between 0 and 2π were assigned to each stimulus onset (black dots), corresponding to its appearance within the cardiac cycle (from the previous to the next R peak in the ECG). Mean (grey arrow) and circular density (grey line) of stimulus onsets were calculated per subject. Individual cardiac phases from the binary analysis (see below) are visualised as circular segments (start: solid, end: dashed): systole (orange) and diastole (blue). For binary analysis (bottom), stimulus onsets (dashed line) were binned into participant-specific systole (orange) and diastole (blue) using a template approach (cf. **Supplementary Methods**, **Supplementary Results**, **Fig. S1, and Fig. S2**).

### Stimuli

The picture set consisted of 180 coloured photographs (60 pictures with positive, 60 with negative, and 60 with neutral content) of humans in various life situations, selected from a collection of standardized and validated affective picture material (EmoPicS) (Wessa et al., 2010). For the index numbers of the selected photographs cf. **Table S1** (**Supplementary Methods**). To correct for potential stimulus-intrinsic influences on visual processing, the three picture sets were largely matched for physical image statistics (Wessa et al., 2010): Contrast and visual complexity did not differ (all p > .21); positive images had significantly higher luminance values than negative images (t(118) = 3.75, p < .001, Cohen’s d = 0.68), while both did not differ from neutral images. In addition, two independent observers matched the three sets for more high-level stimulus features: (1) number of people shown, (2) number of images with social interactions, (3) number of images with close-ups, (4) number of images showing eye contact with the observer. Notably, positive and negative images were matched for (normative) arousal ratings and did not significantly differ (t(78.6) = 1.20, p = .23, Cohen’s d = 0.22). Stimuli were displayed using MATLAB version 7.8.0.347 (The MathWorks Inc., Natick, MA, USA) with the Psychophysics Toolbox 3 (Brainard, 1997; Kleiner et al., 2007; Pelli, 1997).

For each participant, stimuli were randomly selected and presented in randomized order: For encoding, a subset of 120 pictures with 40 pictures of each picture valence (positive, neutral, negative) was sampled from the whole set of 180. The second picture set, shown during the recognition period, consisted of the 60 yet unused pictures (20 per picture valence)—serving as distractors—as well as 60 memory probes (20 per picture valence) that were sampled from the encoded picture set.

### ECG recording

ECG was recorded at 2048 Hz using an ActiveTwo AD amplifier (Biosemi, Amsterdam, Netherlands). Three electrodes were attached according to an adapted limb lead configuration at the right and left lower coastal arch as well as the left medial ankle. Participants were told that the ECG is to measure their general bodily state without mentioning details regarding the experimental conditions (Fiacconi et al., 2016). The ECG lead most clearly displaying the onset of ventricular depolarisation (lead II) was used for analysis.

### Additional measures of inter-individual differences

Previous studies suggest that the influence of cardiac signals on perception and behaviour varies with interoceptive accuracy, that is, the ability to consciously perceive signals originating in the body (Dunn et al., 2010; Garfinkel et al., 2013). We determined inter-individual differences in interoceptive accuracy with a heartbeat perception task (Schandry, 1981), in which participants were asked to estimate the number of their heartbeats in five intervals of different length (25, 45, 15, 55, and 35 s). As inter-individual differences in anxiety have been proposed to moderate the behavioural effect of autonomic signalling (Garfinkel et al., 2014; Pollatos, Schandry, Auer, & Kaufmann, 2007), we also acquired participants’ trait anxiety using the State-Trait Anxiety Inventory (STAI-T; Laux, Glanzmann, Schaffner, & Spielberger, 1981; Spielberger, Gorsuch, Lushene, Vagg, & Jacobs, 1983). Resting heart rate variability (HRV) measures inter-individual differences in brain-heart interaction and particularly in parasympathetic cardioregulation (Task Force, 1996). HRV can be quantified through changes in the beat-to-beat intervals of the ECG (Task Force, 1996). We calculated the root mean square of successive differences (rMSSD) during the 7-minute baseline ECG.

### Procedure

Upon arrival in the lab, participants were equipped with the ECG. Comfortably seated, they were asked to relax for seven minutes and breathe normally. Then, blood pressure was measured (twice, if elevated) using a standard sphygmomanometer (OMRON M8 Comfort). After a brief training session to familiarize participants with the task, they performed the two experimental periods (i.e., encoding and recognition). During a 5-minute break between the two parts, participants completed the Trait Anxiety Inventory (STAI-T) (Laux et al., 1981; Spielberger et al., 1983). To assess subjective perception in our sample and compare it to the EmoPicS’ normative ratings, all 180 photos were rated after the recognition period similarly to the original EmoPicS normative ratings (Wessa et al., 2010): “How do you feel looking at the picture?” was answered for valence (1: sad – 9: happy) and arousal (1: calm – 9: excited) on a 9-level Likert-type scale. For each trial, both rating scales were displayed successively, one above and one below each picture, followed by a 500-ms fixation cross (between trials). Finally, subjects performed the heartbeat perception task (Schandry, 1981) that was presented acoustically.

### Data analysis

The timing of behavioural responses was analysed relative to the heartbeats: electrical events indicating the beginning of each cardiac cycle (R peaks) were extracted from the ECG signal with Kubios 2.2 (Tarvainen, Niskanen, Lipponen, Ranta-aho, & Karjalainen, 2014, http://kubios.uef.fi/). Two complementary analytic approaches—circular and binary analysis—were performed to exploit the oscillatory (repeating cycle of cardiac events) as well as the phasic (two distinct cardiac phases: systole and diastole) nature of cardiac activity, respectively.

### Encoding period—cardiac modulation of self-paced visual sampling

#### Circular analysis

For the circular analysis, we computed the relative onset of each key press (prompting picture onset) within the cardiac cycle, which was indicated in the ECG as the interval between the previous and the following R peak (**Fig. 1b**). According to its relative timing within this R-R interval, radian values between 0 and 2π were assigned to each stimulus (Ohl et al., 2016; Pikovsky, Rosenblum, & Kurths, 2001; Schäfer, Rosenblum, Kurths, & Abel, 1998). For each participant, we computed the mean of the circular distribution for the 120 picture onsets. In a second step, a mean vector of all participants was computed via vector addition of individual means divided by their number, showing the average self-paced picture onset in the cardiac cycle across the group, and weighted by its length (mean resultant length ϱ) to reflect the spread of individual means around the circle. As a measure of concentration of circular data, ϱ was integrated in a subsequent Rayleigh test for uniformity (Pewsey, Neuhäuser, & Ruxton, 2013): if ϱ gets sufficiently high to exceed a threshold value (i.e., the set of individual means is not spread evenly across the cardiac cycle), the data can be interpreted as too locally clustered to be consistent with a uniform distribution that served as null hypothesis (Pewsey et al., 2013). The code for individual and group-level circular analysis can be found on GitHub (https://github.com/SKunzendorf/0303_INCASI). Confidence intervals and significance were non-parametrically calculated through bootstrapping based on analyses from a previous study (Ohl et al., 2016): From the original pool of 43 participants, we drew a random bootstrap sample of 43 participants with replacement. For each participant in the bootstrap sample, we first computed a circular density (bandwidth = 20) of picture onsets, and then computed the mean circular density across the 43 participants in the bootstrap sample. Confidence intervals (95%) were determined as 2.5% and 97.5% percentiles from the distribution of mean circular densities obtained by repeating the bootstrap procedure 10000 times. Deviation from the circular uniform was considered as significant when the 95% confidence interval determined by the bootstrapping is outside the circular density of a uniform distribution.

#### Binary analysis

To account for the phasic nature of cardiac activity and to increase comparability to previous cardiac cycle studies, we segmented the cardiac cycle into systole and diastole (**Fig. 1b**). It needs to be noted that systolic phases vary inversely with heart rate (Fridericia, 1920; Lewis, Rittogers, Froester, & Boudoulas, 1977; Lombard & Cope, 1926; Wallace, Mitchell, Skinner, & Sarnoff, 1963; Weissler, Harris, & Schoenfield, 1968): Although the absolute length of systole decreases with a faster heartbeat, its proportionate share of the entire cardiac cycle increases. Between-subject variation of cardiac phase length (e.g., through differences in heart rate) supports the need to adapt analytical approaches in cardiac cycle studies. With the ECG waveform as physiological reference of cardiac activity, we did not use absolute systole and diastole lengths (e.g., defining systole as the 300 ms following an R peak) but computed participant-specific cardiac phases. Cardiac modulation of perception and cognition has often been attributed to baroreceptor signalling (e.g., Garfinkel & Critchley, 2016; Lacey & Lacey, 1974), which occurs in response to transient pressure rises (i.e., with the systolic upstroke) at each blood ejection (Angell James, 1971). In our approach, phases of high baroafferent feedback were approximated by determining each participant’s systolic ejection phase (in the following referred to as “systole”). For the detailed binning procedure, the time ranges of individualized cardiac phases, and the association between heart rate and cardiac phase length cf. **Supplementary Methods**, **Supplementary Results, Fig. S1, and Fig. S2**. The (self-paced) picture onsets were then assigned to the respective cardiac phase (i.e., individual systole or diastole). To take into account between-subject differences in heart rate (and thus cardiac phase lengths), the sum of picture onsets per phase (as ratio of all 120 trials) was normalized to the proportion of the subject-specific phase length in the total cardiac cycle, resulting in a value of (picture onsets per cardiac phase / 120) / (individual cardiac phase length / individual mean R-R length) for each cardiac phase. With no cardiac effect, button presses (triggering picture onsets) would be randomly distributed across both cardiac phases. That is, the rate of systolic (diastolic) picture onsets should correspond to the proportion of systole (diastole) in the total R-R length, thereby resulting in a ratio of 1. A ratio >1 thus reflects an over-proportional accumulation of picture onsets in the respective cardiac phase. In the group-level analysis, normalized systolic and diastolic ratios were tested against each other with a two-sided paired t-test.

### Recognition period—cardiac modulation of recognition memory

#### Circular analysis

To relate memory performance in the recognition period and stimulus onset in the encoding period, we analysed—for each participant—the stimulus subset of memory probes (i.e., pictures in the recognition period that had already been shown during the encoding phase) with respect to their cardiac onset during encoding. Parallel to the circular analysis regarding visual sampling during encoding (see above), we computed the self-paced onset of memory probes during encoding across the participant’s cardiac cycle (**Fig. 1b**). To correct for a possible bias due to self-paced memory probe distributions, three subjects with non-uniform circular distributions of memory probes (indicated by significant Rayleigh tests) were excluded from further analysis. At the group level, we then analysed circular distributions of onset times for memory probes that were correctly remembered (hits) or erroneously identified as new pictures (misses). To that aim, sets of individual mean onsets for hits and misses were tested against the circular uniform distribution using Rayleigh tests.

#### Binary analysis

The hypothesized association between cardiac phase and memory performance was further analysed by determining the stimulus’ phasic timing during encoding, that is, whether it had been presented in individual systole or diastole (for detailed binning procedure cf. **Supplementary Methods**), relative to its recognition performance. To predict the binary recognition outcome (miss = 0, hit = 1), we computed general linear mixed regression models (GLMM) for binomial data, with subjects as random factor. The first model (m1) was fitted for the overall fixed effect of valence with the three levels of picture valence (positive, negative, neutral) contrast-coded against neutral picture valence as baseline condition. The second model (m2) included picture valence (negative-neutral, positive-neutral), cardiac phase (diastole = 0, systole = 1), and their interaction as fixed effects. Significance was obtained by likelihood ratio tests to compare the full model, which included the effect in question (i.e., valence, cardiac phase), with the reduced model, which did not include the effect in question (i.e., m1 vs. m0 and m2 vs. m1). NB: For m1, the data included only pictures encoded in individual systole or diastole (i.e., pictures encoded in non-defined cardiac intervals were excluded from analysis). An advantage of GLMMs (Jaeger, 2008) is that they can simultaneously test for random effects of subject and item (i.e., picture). To assess additional variance explained by individual pictures, we added picture as a random factor to our models. For the regression analysis, we used the lme4 package (Bates, Mächler, Bolker, & Walker, 2015) in the R Statistical Environment.

### Exploratory analysis of visual sampling and recognition memory relative to inter-individual differences

In an additional exploratory analysis, participants’ individual (i.e., systolic and diastolic) ratios of self-paced picture onsets as well as their overall mean recognition performance (i.e., the percentage of correctly recognised pictures) were correlated with variables of inter-individual differences: Interoceptive accuracy (centred via z-transformation), trait anxiety (centred via z-transformation), and resting heart rate variability/ rMSSD (log-transformed to mitigate skewedness and centred to the mean). For measures of interoceptive accuracy, two participants with lacking information in the heartbeat perception task were excluded from the analysis (n = 41).

### Code availability

The code of our analysis, computed in the R Statistical Environment (v3.4.3) with RStudio version 1.0.136 (RStudio Team, 2016) is available on GitHub (https://github.com/SKunzendorf/0303_INCASI). Graphics were obtained with the circular package (Agostinelli & Lund, 2013) and the ggplot2 package (Wickham, 2009).

### Data availability

The data that support the findings of this study are available on GitHub (https://github.com/SKunzendorf/0303_INCASI). Data for preprocessing are available from the corresponding author upon request.

## Results

We tested whether self-paced visual sampling and visual memory encoding fluctuate across the cardiac cycle. Based on previous studies that demonstrated facilitated visual processing (Pramme et al., 2014, 2016) and increased oculomotor activity (Ohl et al., 2016) during the early phase of the cardiac cycle, we hypothesized that participants prefer to prompt a visual stimulus during early phases of the cardiac cycle. In addition, we hypothesized memory performance to be influenced by the cardiac time point of memory probes during encoding (Garfinkel et al., 2013).

### Encoding

The distribution of self-paced picture onsets relative to the cardiac R-R interval showed an overall increase in early phases of the cardiac cycle (M = 0.33π, SD = 0.52π, ϱ = 0.26, cf. **Fig. 2a**). This observation was supported by inferential circular statistics indicating a non-significant trend that the self-paced key presses in the present experiment are unlikely to be uniformly distributed (Rayleigh test statistics R_0_ = 0.26, p = .053). Of note, the same analysis with our preregistered sample size (i.e., the first 40 healthy subjects of N = 43; cf. https://osf.io/5z8rx/) showed a significant deviation from a uniform distribution (R_0_ = 0.28, p = .039), that is due to a significant increase in self-paced key presses in the interval from 0.24 to 0.44π (**Fig. 2a**) as revealed by nonparametric bootstrapping (performed on the original participant pool, N = 43). Although this circular statistics can infer that the distribution of relative picture onsets deviates from a uniform distribution, it cannot pinpoint the transition from systole to diastole – particularly in the presence of varying heartbeat lengths, for which the same section of the circular distribution can be associated with different cardiac phases.

**Figure 2.**
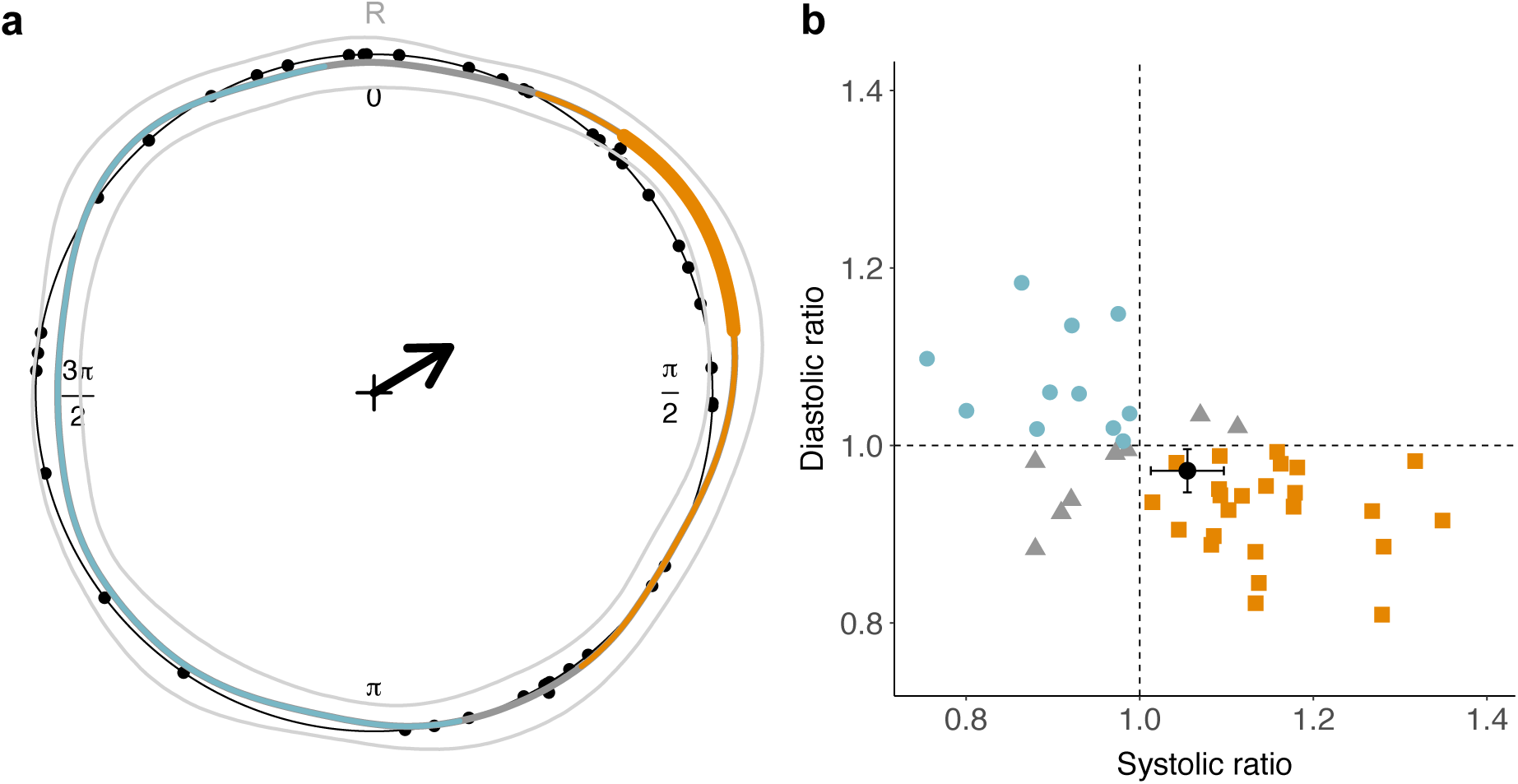
Circular and binary analysis of visual sampling relative to the heartbeat. ***a***, Circular distribution of individual mean picture onsets (black dots, N = 43) across the cardiac cycle (from R peak to R peak). We observed a trend for increased self-prompted stimulus presentations (weighted overall mean as black arrow) in early phases of the cardiac cycle. Based on a bootstrapping procedure, we computed the mean circular density of picture onsets (middle thicker line), as well as a 95% confidence interval (CI, within inner and outer thin grey lines). Segments of the cardiac cycle are determined as statistically significant (thick orange segment) when the circular density significantly differs from the circular uniform (i.e., the lower bound of the CI is outside of the black uniform circle). To relate segments of the cardiac cycle to the two cardiac phases (systole = orange, diastole = blue, non-defined = grey), overall mean systole and diastole lengths were obtained, showing, that the significant density segment falls into systole. ***b***, Most subjects (24/43) preferred to prompt pictures during their systole (orange/square). Fewer subjects (11/43) chose to prompt them during their diastole (blue/dot) or did not show a preference in any of the two defined phases (8/43; grey/triangle). The phase-specific proportion of key presses (relative to all 120 trials) was normalized by the proportion of the cardiac phase (systole, diastole) in the entire R-R interval. A ratio >1 thus indicates that the number of prompted picture onsets during systole or diastole exceeds the number that would be expected if they were uniformly distributed (resulting in a ratio = 1, dashed line). The group-level mean (black dot with standard error bars) shows an over-proportional accumulation of picture onsets during individual systole relative to individual diastole.

Accounting for the bi-phasic nature of cardiac activity in the binary analysis, we computed the number of key presses during cardiac systole and diastole, normalised by the proportion of the systole vs. diastole in the whole cardiac cycle. We found a significantly larger (t(42) = 2.76, p = .009, Cohen’s d = 0.42) ratio of picture onsets in the systole (M = 1.05, SD = 0.14) as compared to diastole (M = 0.97, SD = 0.081), corroborating our finding of an increase in self-paced visual sampling during cardiac systole (cf. **Fig. 2b**). Similar results were obtained for the preregistered sample size (n = 40), showing a significantly larger (t(39) = 2.70, p = .010, Cohen’s d = 0.43) systolic (M = 1.05, SD = 0.14) than diastolic ratio (M = 0.97, SD = 0.079).

### Recognition

We also investigated the association between the cardiac cycle and memory processing: In a circular analysis, we tested whether the distribution of onset times during stimulus encoding differed for pictures that were correctly remembered (hits) or erroneously identified as new pictures (misses). Overall, picture onset times during the encoding period did not significantly deviate from a uniform distribution over the cardiac cycle for hits (R_0_ = 0.20, p = .21) and misses (R_0_ = 0.16, p = .36).

We further investigated the influence of cardiac phase (systole, diastole) and picture valence on recognition memory. The results of the GLMM (cf. **Table 1**) with contrast-coded picture valence (m1), that is, negative-neutral and positive-neutral, showed a significant memory benefit for negative vs. neutral and for positive vs. neutral stimuli. More specifically, memory performance for negative pictures (M = 0.80, SD = 0.13) and for positive pictures (M = 0.78, SD = 0.16) significantly exceeded memory performance for neutral pictures (M = 0.73, SD = 0.18). Comparison of m1 against the null model (without the fixed effect of valence) showed that valence significantly increased the model fit. Critically, adding cardiac phase (i.e., systole, diastole) to the model (m2) did not improve the model fit (cf. **Table 1**): Neither phase nor its interaction with picture valence significantly accounted for variation in recognition memory. Compared to m1, cardiac phase did not significantly improve the model fit. Adding picture as additional random effect only slightly changed parameter estimates but, critically, did not account for additional variance in the association between memory performance and the cardiac cycle (detailed results not reported).

**Table 1.**
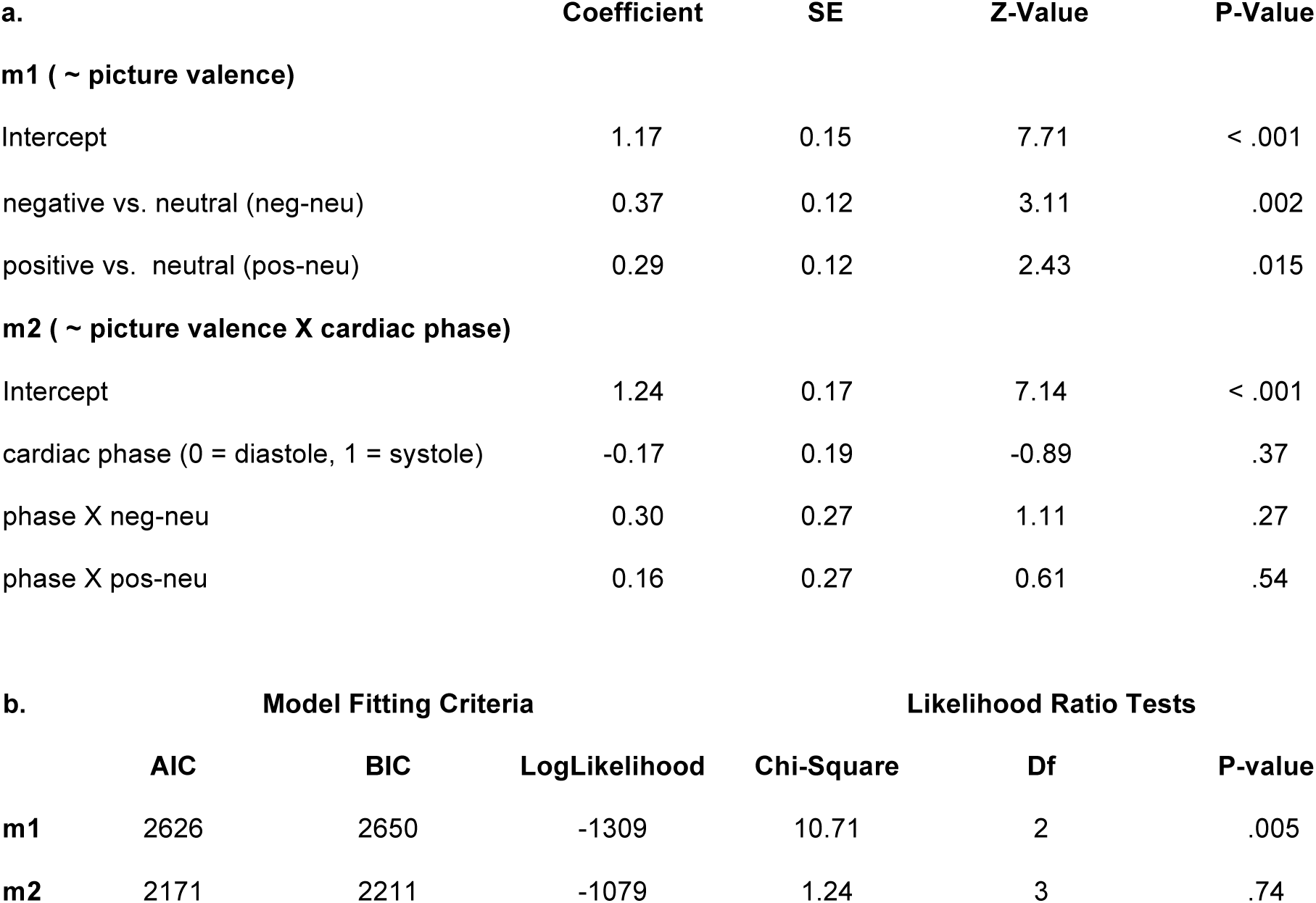
***a***, General linear mixed model (GLMM) with recognition memory (hit = 1, miss = 0) relative to picture valence (negative-neutral, positive-neutral) (m1 = memory ~ picture valence + (1|vp)), cardiac phase, and their interaction (m2 = memory ~ picture valence × cardiac phase + (1|vp)). Valenced pictures (negative and positive) showed a significant memory benefit compared to neutral pictures. Neither cardiac phase nor its interaction with picture valence significantly accounted for variation in visual memory performance. ***b***, Likelihood ratio tests of m1 and m2 against the reduced model (i.e., m0 and m1, respectively) show that picture valence significantly increased the model fit while cardiac phase did not account for variation in memory performance.

| a. | Coefficient | SE | Z-Value | P-Value |
| --- | --- | --- | --- | --- |
| **m1 ( ~ picture valence)** |  |  |  |  |
| Intercept | 1.17 | 0.15 | 7.71 | < .001 |
| negative vs. neutral (neg-neu) | 0.37 | 0.12 | 3.11 | .002 |
| positive vs. neutral (pos-neu) | 0.29 | 0.12 | 2.43 | .015 |
| **m2 ( ~ picture valence X cardiac phase)** |  |  |  |  |
| Intercept | 1.24 | 0.17 | 7.14 | < .001 |
| cardiac phase (0 = diastole, 1 = systole) | -0.17 | 0.19 | -0.89 | .37 |
| phase X neg-neu | 0.30 | 0.27 | 1.11 | .27 |
| phase X pos-neu | 0.16 | 0.27 | 0.61 | .54 |

**Table 1. a**, General linear mixed model (GLMM) with recognition memory (hit = 1, miss = 0) relative to picture valence (negative-neutral, positive-neutral) (m1 = memory ~ picture valence + (1|vp)), cardiac phase, and their interaction (m2 = memory ~ picture valence X cardiac phase + (1|vp)).
| **b.** | **Model Fitting Criteria** |  |  | **Likelihood Ratio Tests** |  |  |
| --- | --- | --- | --- | --- | --- | --- |
|  | **AIC** | **BIC** | **LogLikelihood** | **Chi-Square** | **Df** | **P-value** |
| **m1** | 2626 | 2650 | -1309 | 10.71 | 2 | .005 |
| **m2** | 2171 | 2211 | -1079 | 1.24 | 3 | .74 |

Beyond our preregistered hypotheses, additional results were obtained by further exploratory analyses.

### Systole-associated visual sampling and inter-individual differences

Individual systolic ratios of self-paced picture onsets were neither significantly correlated with inter-individual differences in interoceptive accuracy (r(39) = -.16, p = .32) nor in heart rate variability (i.e., resting rMSSD; r(41) = .064, p = .69). There was a non-significant (and not hypothesized) trend (**Fig. 3b**) for individual systolic ratios to increase with higher trait anxiety (r(41) = .29, p = .062). Neither interoceptive accuracy (r(39) = .25, p = .11) nor trait anxiety (r(41) = -.21, p = .17) nor heart rate variability (r(41) = .16, p = .32) significantly modulated individual diastolic ratios of picture onsets (**Fig. 3**).

**Figure 3.**
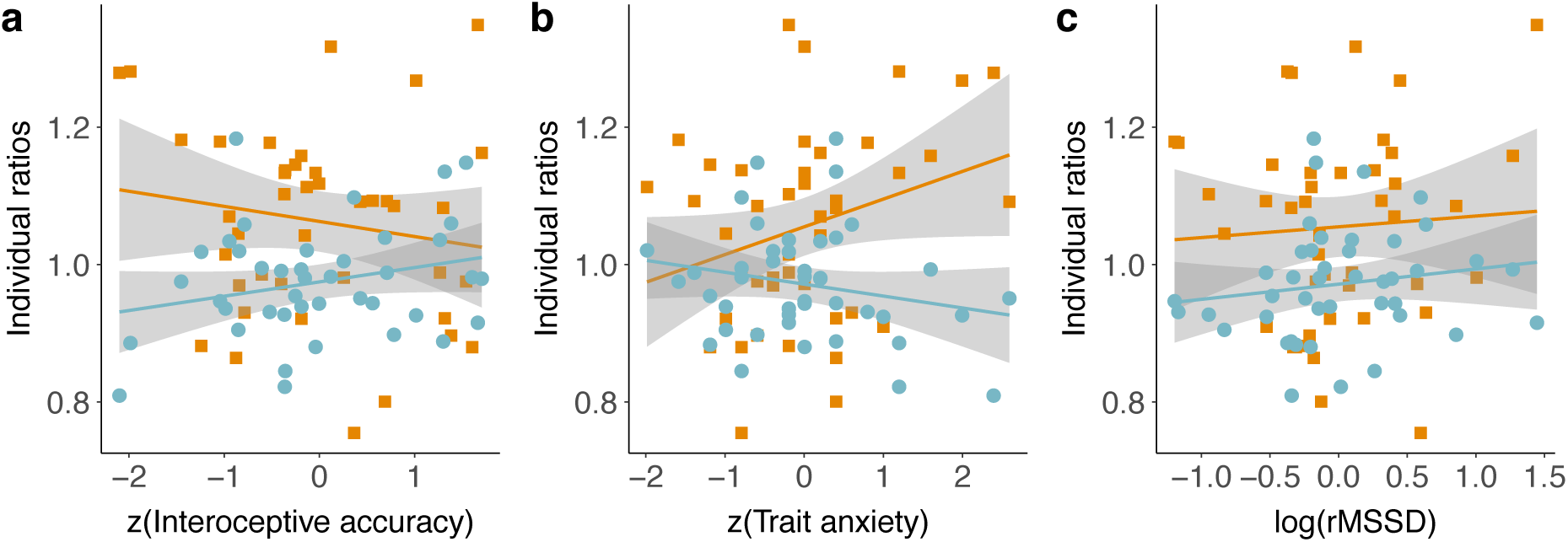
**Correlation of systolic (orange squares) and diastolic (blue circles) ratios of picture onsets with inter-individual differences** in ***a***, Interoceptive accuracy, ***b***, trait anxiety, and ***c***, resting heart rate variability (root mean square of successive differences, rMSSD). Grey areas are the 95% confidence intervals of the respective linear models (orange / blue lines).

### Recognition memory varies and inter-individual differences

We furthermore investigated the role of inter-individual variables (i.e., interoceptive accuracy, trait anxiety, resting heart rate variability) for memory performance with correlation analyses. Neither differences in interoceptive accuracy (r(39) = -.12, p = .46), nor in trait anxiety (r(41) = .039, p = .80) were associated with mean recognition performance (**Fig. 4**). However, there was a non-significant trend of resting heart rate variability (i.e., resting rMSSD) to be positively correlated with mean recognition performance (r(41) = .29, p = .056), that is, recognition memory increased with higher resting heart rate variability (**Fig. 4c**).

**Figure 4.**
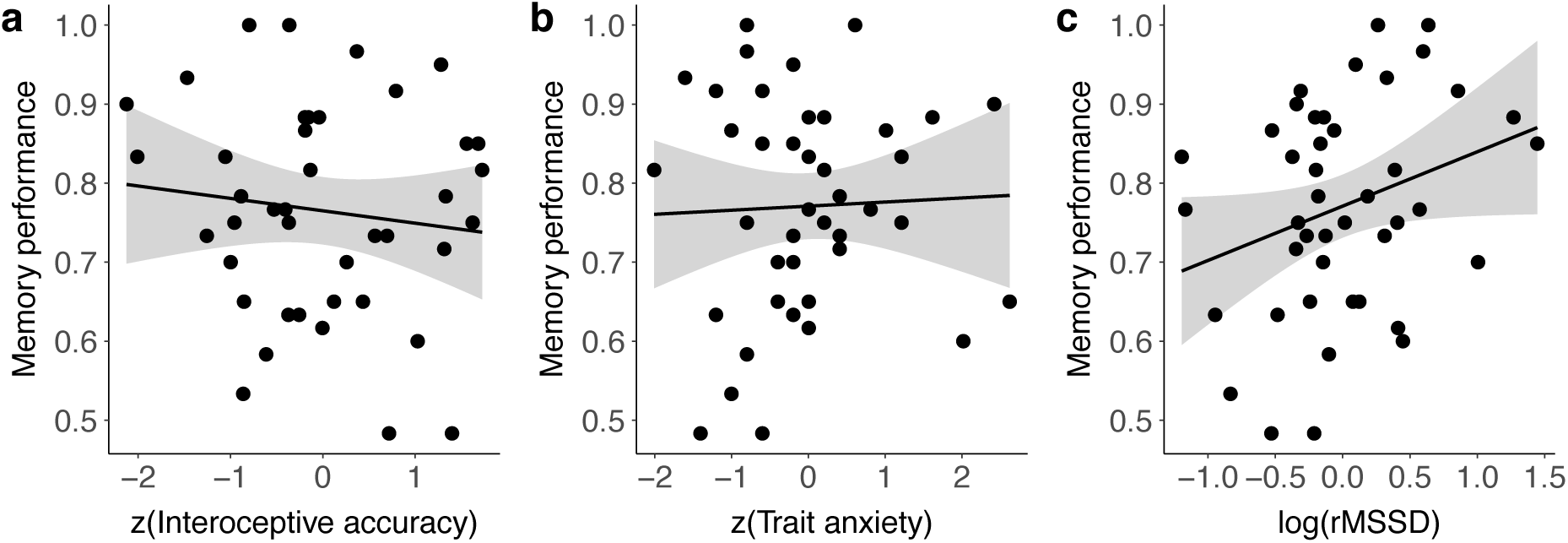
Correlation of mean recognition performance with inter-individual differences. ***a***, interoceptive accuracy, ***b***, trait anxiety, and ***c***, resting heart rate variability (root mean square of successive differences, rMSSD). Grey areas are the 95% confidence intervals of the linear model (black line).

### Subjective perception of picture emotionality

As a control, we analysed if the subjective valence and arousal ratings in our study differed from the EmoPicS normative ratings (**Fig. 5**). For ratings of the two affective dimensions, valence and arousal, mixed-design ANOVAs tested the main and interaction effects of the repeated-measures factor rating category (normative, individual) and the factor picture valence (positive, neutral, negative). We observed a significant main effect of rating category for both, valence and arousal: Mean individual valence ratings (M = 5.06, SD = 1.43) were significantly lower (F(1,352) = 10.1, p = .002) than normative ratings (M = 5.23, SD = 1.69), as were mean individual arousal ratings (M = 3.85, SD = 1.09) compared to normative (M = 4.37, SD = 1.28) ratings (F(1,352) = 27.0, p < .001). However, while valence ratings did not show a significant interaction for rating category × picture valence (F(2,352) = 1.91, p = .15; see **Fig. 5a**), the difference between individual and normative arousal ratings was influenced by picture valence (F(2,352) = 5.71, p = .004; see **Fig. 5b**). To further examine this interaction, two-sided paired t-tests were calculated and p-values were adjusted for multiple comparisons with Bonferroni correction: positive (individual: M=4.04, SD=0.65; normative: M=4.90, SD=0.58) arousal ratings differed significantly larger from each other (t(59) = 20.2, p = < .001, Cohen’s d = 1.39) than both neutral (individual: M=2.91, SD=0.34; normative: M=3.09, SD=0.32) (t(59) = 4.77, p = < .001, Cohen’s d = 0.52) and negative (individual: M=4.60, SD=1.26; normative: M=5.13, SD=1.41) arousal ratings (t(59) = 9.32, p = < .001, Cohen’s d = 0.40). Furthermore, while normative arousal ratings did not differ significantly between positive and negative pictures (cf. **Methods**), individual arousal ratings were significantly higher for negative compared to positive pictures (t(88.3) = 3.05, p = .003, Cohen’s d = 0.56).

**Figure 5.**
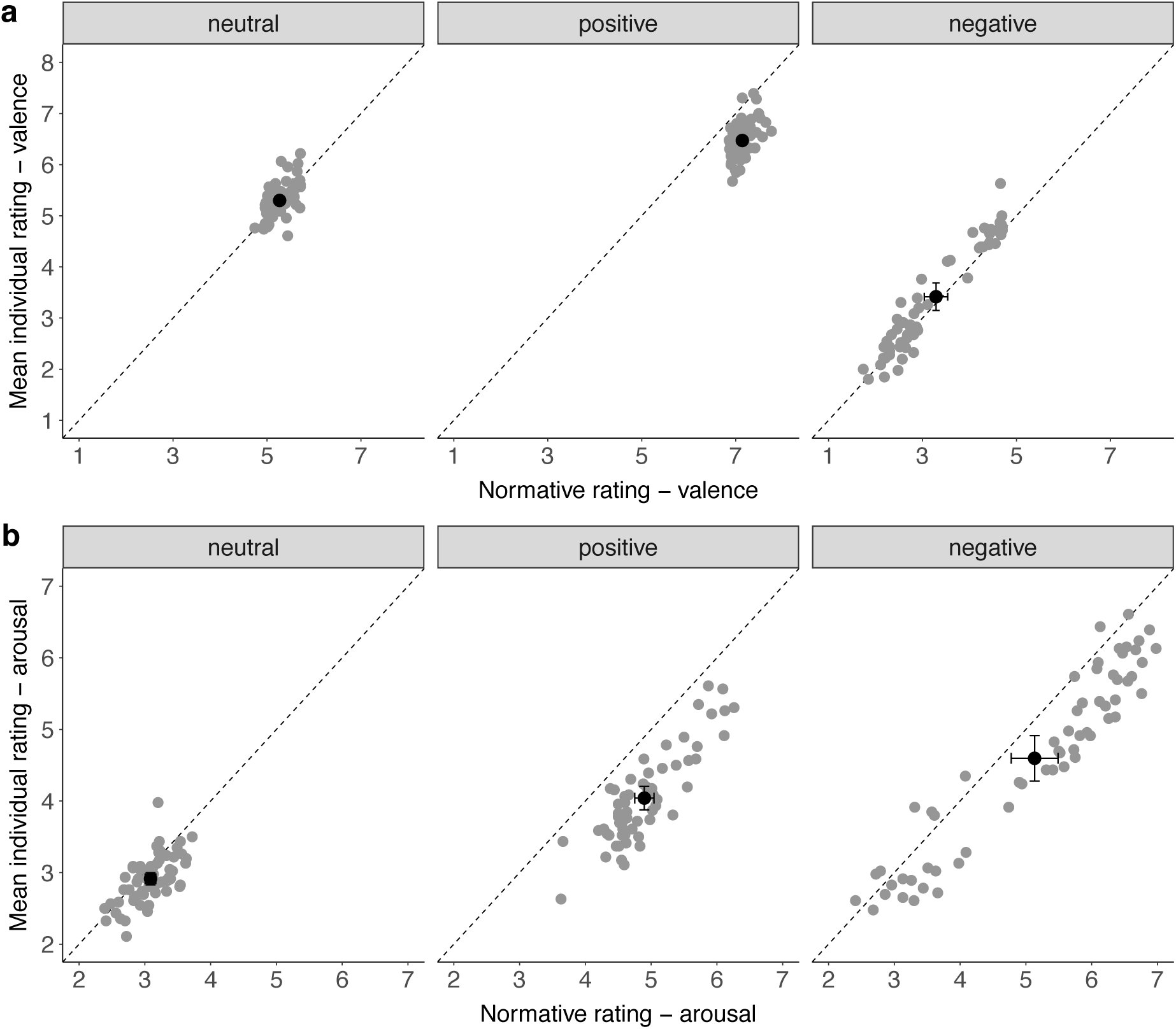
Subjective perception of picture emotionality (individual valence and arousal ratings) compared to normative picture ratings. ***a***, *Valence ratings* and ***b**, Arousal ratings* were both significantly lower for individual compared to normative ratings (overall mean in black with standard error bars). For arousal but not for valence ratings there was a significant interaction effect between rating category and picture valence.

## Discussion

We studied the association between the cardiac cycle and self-paced visual sampling as well as visual recognition memory for pictures of different emotional valence. We hypothesized that facilitated visual processing (Pramme et al., 2014, 2016)—observed specifically for relevant or emotionally salient stimuli (Azevedo et al., 2018, 2017; Garfinkel et al., 2014)—as well as facilitated oculomotor processing during systole (Ohl et al., 2016) guides active perception in the shape of a preference to prompt a relevant visual stimulus during early phases of the cardiac cycle. We observed a significant accumulation of key presses (i.e., prompted picture onsets) during systole, thereby showing for the first time a coupling between self-paced visual sampling and the heartbeat. Memory performance, however, was only influenced by picture valence, replicating a significant memory benefit for emotional content (Hamann, 2001; Kensinger, 2006; Kensinger & Corkin, 2003; LaBar & Cabeza, 2006), but not further modulated by the cardiac phase in which targets were encoded (Garfinkel et al., 2013). The association between the cardiac cycle and self-initiated actions complements findings of facilitated visual processing during systole, thereby proposing a link between the heartbeat and active perception.

Sensorimotor processing of passively presented stimuli has been shown to be decreased during early cardiac phases (Birren et al., 1963; Callaway & Layne, 1964; Edwards et al., 2007; Lacey & Lacey, 1974; McIntyre et al., 2008; Saari & Pappas, 1976), but evidence is mixed, including some earlier null findings (Jennings & Wood, 1977; Salzman & Jaques, 1976; Thompson & Botwinick, 1970). However, which processing stages are modulated by the heartbeat has long remained unclear. More fine-graded decomposition of reaction time into central (sensory, pre-motor) and peripheral (motor) processes has indicated that the response inhibition during early cardiac phases was confined to central (pre-motor) levels of stimulus processing, whereas the motor component remained unaffected (Edwards et al., 2007; McIntyre et al., 2008; Saari & Pappas, 1976) or even accelerated (Schulz et al., 2009). This differential effect could also underlie findings that report an increased tendency to act during systole such as fire a virtual (Azevedo et al., 2017) or an actual weapon (Mets et al., 2007) during early cardiac phases. Our results of facilitated spontaneous actions during systole when engaging with a visual stimulus furthermore add to results from oculomotor behaviour: Ohl et al. (2016) reported involuntarily occurring oculomotor activity (i.e., microsaccade generation) to be heightened during early cardiac phases. Hence, for both somaomotor and oculomotor processes, our readiness to act upon an external stimulus fluctuates with our (internal) cardiac rhythm, being relatively increased during the phase of systolic blood ejection.

Although underlying heart-brain pathways remain unclear, systolic influences on perception and cognition have often been attributed to the phasic nature of cardio-afferent signalling, which is triggered with the heartbeat (Koriath & Lindholm, 1986; Lacey & Lacey, 1974, 1978). More specifically, stretch-responsive baroreceptors located in arterial walls respond to transient pressure rises at each blood ejection and communicate the current cardiovascular state (i.e., the heartbeat’s strength and timing) to the brain. Thus, baroreceptor-transmitted cardiac signals, which are phasically registered by central processing systems, have been proposed to induce a general suppression of cortical excitability, converging with earlier findings of sensory inhibition (Dembowsky & Seller, 1995; Rau & Elbert, 2001). Formulated in terms of the *interoceptive predictive coding* framework (Barrett & Simmons, 2015; Seth, 2013), which extends the *free-energy principle* (Friston, 2010) to interoceptive processes and connects them to consciousness and emotion, this periodic—and thus predictable—ascending baroreceptor input to central structures is anticipatorily cancelled out by top-down interoceptive predictions, thereby minimizing its influence on perception (Critchley & Garfinkel, 2018; Salomon et al., 2016). Besides modulating arterial baroreceptor discharge, heartbeat-related pressure fluctuations generate periodic sensory influences throughout the body—affecting for example the discharge of tactile (Macefield, 2003) or muscle spindle afferents (Birznieks, Boonstra, & Macefield, 2012)—which are predicted and normally do not enter perceptual awareness. In our constant attempt to minimize sensory uncertainty (Peters, McEwen, & Friston, 2017), inhibition of such predictable cardiac-induced sensory effects has been argued to reduce—potentially distracting—self-related sensory noise (Salomon et al., 2016) at the benefit of our processing of the outside world, for example by increasing the signal-to-noise ratio of external stimuli. Correspondingly, stimuli presented simultaneously with the heartbeat are interpreted as sensory consequences of the organism’s own (internal) cardiac activity and thus perceptually attenuated (Salomon et al., 2016). Our finding that participants implicitly act upon a visual stimulus during the phase of heartbeat-related (baroreceptor-mediated) central inhibition could indicate a short-term benefit: As argued by Pramme et al. (2016), the impact of baroreceptor-mediated central influences may depend on context- or task-specific processing demands. In other words, extraction of behaviourally relevant external stimuli from a distracting sensory scene might be facilitated during inhibition of irrelevant (e.g., cardiac-related) sensory information (Pramme et al., 2016). Accordingly, predictable phases of attenuated heartbeat-related noise might provide a short-term window to facilitate active engagement towards an external relevant stimulus.

Such differential processing during transient cardiac signalling converges with the observed specificity of cardiac effects when using valenced stimuli, in particular selectively facilitated processing of threat stimuli during systole (Garfinkel & Critchley, 2016). The notion that cardiac signals prioritise the processing of motivationally relevant information suggests a crucial role of cardiac interoceptive information in conveying bodily arousal states to the brain (Critchley & Garfinkel, 2018; Garfinkel & Critchley, 2016). In other studies, states of higher psychophysiological arousal have shown to bias the processing of relevant stimuli, including facilitated memory formation (Cahill & McGaugh, 1998; Mather et al., 2016; Mather & Sutherland, 2011; McGaugh, 2015). However, the pattern of results concerning memory modulation across the cardiac cycle remains fragmented and unclear. Although perceptual sensitivity for emotional stimuli is increased during systole (Garfinkel et al., 2014), affective (positive, negative) and neutral words are recalled less often when they are encoded during systole as compared to diastole (Garfinkel et al., 2013). Furthermore, this cardiac memory effect could only be obtained in subjects with lower interoceptive accuracy, suggesting influences of an individual’s access to interoceptive sensations (Pollatos & Schandry, 2008). On the other hand, Fiacconi et al. (2016) found that fearful and neutral faces presented during systole are more likely to be judged as known or old, irrespective of whether they had been shown before or not. In our study, recognition memory was not influenced by the cardiac phase during which a stimulus was encoded—also adding to a recent study reporting a lack of cardiac influences on memory retrieval (Pfeifer et al., 2017)—but only by its valence, showing a significant benefit for negative and positive pictures. This suggests that—at least in our study—the influence of externally-induced emotional arousal states (e.g., by seeing an upsetting negative picture) on memory formation (Tambini et al., 2017) might have exceeded a transient cognitive modulation across the cardiac cycle (Garfinkel et al., 2013). Such reasoning is further supported by a recent study showing that interoceptive cardiac signals can easily be overshadowed by external stimuli or other task-specific influences (Yang, Jennings, & Friedman, 2017). Although in our study, stimulus content was largely matched along several dimensions (physical image statistics and more high-level features), differences in stimulus features may still account for variation in memory effects associated with the cardiac cycle. For example, differences have been reported for different stimulus categories like words (Garfinkel et al., 2013), faces (Fiacconi et al., 2016), and complex scenes (present study), but also for low-level stimulus properties such as spatial frequency (Azevedo et al., 2018). However, accounting for picture as random effect in our GLMM analyses did not explain additional variance in memory performance across the cardiac cycle.

Our exploratory supplementary finding indicates that inter-individual differences in recognition memory are positively associated with inter-individual differences in resting heart rate variability. Considered a trait marker of autonomic or parasympathetic cardio-regulation and—more generally—of heart-brain coupling (Thayer et al., 2012), variation in beat-to-beat intervals at rest has been associated with cognitive capacities: participants with higher resting heart rate variability performed better in tests of working memory and attention (Hansen, Johnsen, & Thayer, 2003; Luft, Takase, & Darby, 2009). Future studies investigating cardiac influences on cognition could further examine the impact of inter-individual differences in resting heart rate variability.

Taken together, cardiac phase effects on perception, cognition, and behaviour might constitute a non-functional epiphenomenon emerging from transient physiological changes, which set the context for heart-brain interactions: As baroreceptor-transmitted afferent signals (Critchley & Harrison, 2013) occur with every systolic pressure wave and reflect momentary states of increased blood pressure, they constitute a fine-tuned reference of cardiovascular arousal. Although autonomously generated, cardiac fluctuations are integrated in multiple feedback loops to react to environmental challenges such as exercise, body position, or stress (Dampney et al., 2002; Glass, 2001; Saper, 2002). Cardiovascular arousal is thus directly encoded via frequency (e.g., increased heart rate) and waveform (e.g., increased amplitude in elevated blood pressure) (Dampney et al., 2002; Schächinger, Weinbacher, Kiss, Ritz, & Langewitz, 2001), which reciprocally affect afferent baroreceptor stimulation (Chapleau, Li, Meyrelles, Ma, & Abboud, 2001). Influences of the cardiovascular state on sensory processing might thus subtly emerge with heartbeat-related pressure fluctuations, but are only fully expressed under a sustained shift of our bodily state beyond physiological variability; for example, under stress, when the whole spectrum of adaptive brain-body responses is activated (e.g., elevated blood pressure and accelerated heart rate) and the afferent cardiac signalling increases. Correspondingly, Luft and Bhattacharya (2015) found that the representation of cardiac signals in the brain, as measured by EEG-derived heartbeat-evoked potentials, differs between states of high vs. low emotional arousal. Besides increased amplitudes in arterial pressure waves, it could be argued that a faster heartbeat during stressful situations—next to providing metabolic support for action requirements to restabilize our homeostatic integrity (Gianaros & Wager, 2015)—results in relatively increased systolic signalling (as raises in heart rate occur mainly at the expense of diastole length) and thereby generates more time windows of selectively facilitated sensory processing. Evidence for this proposal comes from a recent study that associated experimentally increased heart rates (Pezzulo et al., 2018) with prioritized fear processing across different measures (reaction time, peak velocity, response acceleration, choice uncertainty). Hence, increased signalling of cardiovascular states under conditions of higher bodily arousal and heart rates might more strongly modulate cognition and behaviour (e.g., active perception and self-paced action), thereby supporting what information is preferentially processed.

Our experiment has several limitations: While linking active perception and the cardiac cycle, our design does not allow to decompose cardiac influences at the levels of somatomotor, sensory, and cognitive processing. A control condition to dissociate cardiac-related motor activity from visual processing could rule out a pure motor effect, for example by testing spontaneous motor actions that are not explicitly coupled to perception of (relevant) stimuli. The hypothesis would be that button presses that do not prompt relevant sensory input would be randomly (i.e., uniformly) distributed across the cardiac cycle. An essential step to investigate the role of cardiac activity as bodily reference would be to test cardiac coupling of stimulus processing under conditions of altered cardiac activity (e.g., increased heart rate), for example by inducing stress through increased sensory uncertainty (e.g., by manipulating stimulus predictability). In addition, measurements of cardiac representations in the brain (e.g., using EEG) would extend our understanding of the central integration and modulation of cardiac signals, for example, in the context of self-paced visual sampling and visual memory processing.

In conclusion, our findings imply that the heartbeat constitutes a crucial bodily signal that is integrated in our active engagement with the external world. Specifically, they suggest that we tend to act in a phase of inhibited cardiac- and thus self-related sensory processing (namely cardiac systole) when extracting relevant information from our environment. Subtly emerging under normal conditions, this influence might become functionally relevant in states of high arousal (e.g., in stressful situations). Extending previous frameworks of mind-brain-body interactions (Park & Tallon-Baudry, 2014), we propose that we implicitly exploit internal ongoing bodily fluctuations as a predictable reference frame from which interaction with our ever-changing and often unpredictable environment can arise. When initiating actions to sample the noisy world around us, we relate them to the rhythm we know best—our own heartbeat.

