## Supplementary Methods for "Active information sampling varies across the cardiac cycle"

1. Supplementary Method.

1.1 Index numbers of all EmoPicS selected for our experiment.

| | |
|---|---|
| **Positive** | 1–6, 8–11, 13, 15, 17–18, 20–34, 36, 38–40, 42–57, 61–63, 66, 69, 73–78 |
| **Neutral** | 79-88, 92, 94, 99, 101, 103, 106, 109, 111, 114, 119, 121, 123, 126, 130–132, 135, 138, 142, 145–148, 150, 153, 157–158, 164, 166, 168–172, 174–184, 187, 193, 199, 203–204 |
| **Negative** | 89, 91, 95, 107, 110, 112–113, 116, 118, 125, 127, 139, 141, 144, 159, 161, 167, 189 –190, 202, 207–232, 234–235, 238, 241, 245–247, 249–255 |

**Table S1.** EmoPicS (Wessa et al., 2010) index numbers of the selected stimuli (60 positive, 60 neutral, and 60 negative).

1.2 Detection of individual cardiac phases. Subject-specific systole and diastole lengths were computed based on cardio-mechanical events that could be related to the ECG trace. Although mechanical systole cannot be automatically derived from the duration of electrical systole (Fridericia, 1920), both are closely linked under normal conditions (Boudoulas, Geleris, Lewis, & Rittgers, 1981; Coblentz, Harvey, Ferrer, Cournand, & Richards, 1949; Fozzard, 1977; Fridericia, 1920; Gill & Hoffmann, 2010; T. Lewis, 1920; Wiggers, 1921, 1923). That is, systolic contraction of the ventricles is preceded by their depolarisation (indicated in the ECG by the QRS complex), while aortic valve closure, marking the end of systolic blood outflow, coincides with ventricular repolarisation around the end of the T wave (Gill & Hoffmann, 2010). Based on the ECG, the total ventricular systolic phase (cf. **Fig. S1a**) is described as the time difference between Q wave onset and the T wave end (QT) (e.g., Fridericia, 1920), comprising systolic intervals of depolarization and isovolumetric contraction (no blood outflow, PEP), as well as ejection (blood outflow into the aorta and pulmonary artery, EP). Diastole, during which the ventricles relax and get refilled with blood, describes the remaining part of the cardiac cycle until the onset of the subsequent QRS complex (Boudoulas, Rittgers, Lewis, Leier, & Weissler, 1979; Levick, 2010; R. P. Lewis, Rittogers, Froester, & Boudoulas, 1977; Pappano & Wier, 2013).

As our phase of interest (i.e., baroreceptor activity) is defined by the time course of the pulse pressure wave at each heartbeat (Angell James, 1971; Coleridge, Coleridge, Poore, Roberts, & Schultz, 1984; Levick, 2010), it approximately coincides with the EP as systolic pressure waves activate aortic and carotid baroreceptors within 10–15 ms (i.e., R + 90 ms) and 40–65 ms (i.e., R + 140 ms) after blood ejection, respectively (Rushmer, 1976, cited in Edwards, Ring, McIntyre, & Carroll, 2001; Edwards, Ring, McIntyre, Winer, & Martin, 2009; Quelhas Martins, McIntyre, & Ring, 2014). For each cardiac cycle, the EP was extracted by removing the PEP from the total electrical systole (i.e., QT interval), using regression equations (Weissler, Harris, & Schoenfeld, 1969; Weissler, Harris, & Schoenfield, 1968) that were formerly used in clinical cardiology as a “noninvasive” technique to determine systolic time intervals (R. P. Lewis et al., 1977). Thus, participant-specific systole templates (with an individualized systole length per participant) started approximately with the opening of the aortic valve initiating the blood outflow shortly after the R peak (Weissler, Harris, & Schoenfeld, 1969) and finished with the T wave end, around which the aortic valves close (Gill & Hoffmann, 2010). Individual diastolic phases, which differed (within-subject) from trial to trial due to beat-to-beat heart rate variability, started after a buffer interval of 50 ms and extended to the onset of the Q wave from the following QRS complex as beginning of ventricular depolarization (i.e., diastole was calculated as RR–QT–50 ms).

Through excision of the PEP before the EP as physiologically distinguishable interval of ventricular depolarization and contraction as well as a 50-ms window after the EP, we wanted to avoid a possible overlap between the cardiac phases; with the aim to compare the two distinct cardiac intervals of alternating baroreceptor activity (cf. **Fig. S2**): the systolic phase of blood ejection (activated baroreceptors) vs. the diastolic phase of ventricular relaxation and filling (quiescent baroreceptors).

T wave end detection followed a two-step procedure: First, a T wave template was computed for every participant by averaging 1000 ECG trace snippets from the experimental period, which encompassed a—physiologically plausible—time interval to contain the T wave: up to 390 ms following each R peak. Subsequently, the Trapez area algorithm

(Vázquez-Seisdedos et al., 2011) was applied to compute the T wave end in each subject-specific template: Having located the T peak as local maximum within the template, the algorithm computes a series of trapezes along the descending part of the T wave signal, defining the T wave end as the point where the trapezium's area gets maximal.

Q wave onsets (or, if not clearly discernable, R peak onsets), demarcating the onset of ventricular systole, served as a starting point for the regression equations (Weissler et al., 1968). Similarly computed within an averaged template, they were determined as the first prominent negative deflection from baseline (or, if replaced by R peak onset, the first dominant upward deflection), preceding the R peak in the ECG trace (Sherwood et al., 1990). Thereby, a template-based systole length (QT) was obtained for each participant.

To exclude PEP intervals from participants' total systole length (QT), a regression equation was applied to each cardiac cycle, which relates the duration of PEP to the participant's mean heart rate (HR), and is corrected for slight differences between male (M) and female (F) (PEP (M) = -0.4 HR + 131, PEP (F) = -0.4 HR+ 133) (R. P. Lewis et al., 1977; Weissler et al., 1969, 1968). The PEP (i.e., Q wave onset to the approximated opening of the aortic wall, obtained by measuring the beginning systolic upstroke) was then subtracted from the whole QT interval, which determined the EP within each cardiac cycle. We visually checked the fit of individual cardiac intervals, visualizing them on the actual ECG trace (in the encoding period) of each subject (cf. **Fig. S1a)**. The code for detection of individual cardiac intervals (in R) is available on GitHub (https://github.com/SKunzendorf/0303_INCASI).

### 2. Supplementary Results.

**2.1 Inter-individual variation of cardiac intervals.** To illustrate the heart rate-dependent variation of cardiac intervals, the two tachycardic subjects were included in this control analysis (sample size: $N$ = 45, including 21 female; age: 18–34 years, $M$ = 25.9 years, $SD$ = 4.38). Mean heart rates varied from 52.6 to 113.7 bpm ($M$ = 74.7, $SD$ = 12.9), with mean RR intervals ranging from 0.53 s to 1.15 s ($M$ = 0.83, $SD$ = 0.14).

Visualization of subject-specific systolic templates within their individual ECG trace indicated that under the present experimental conditions, within-subject systole lengths stay rather constant. However, between subjects, systolic intervals do differ together with differences in mean individual heart rates.

In line with earlier research (Boudoulas et al., 1981; R. P. Lewis et al., 1977; Lombard & Cope, 1926; Wallace, Mitchell, Skinner, & Sarnoff, 1963; Weissler et al., 1969, 1968), we observed an inverse correlation of EP length and mean individual heart rate (Pearson's $r(43) = -0.72$, $p < 0.001$), confirming that heart rate is an important determinant of EP. As shown in **Fig. S1b left**, absolute EP duration decreased with higher resting heart rates, ranging from 0.23 s to 0.31 s ($M$ = 0.26, $SD$ = 0.020), with a total systolic QT interval between 0.32 s and 0.42 s ($M$ = 0.36, $SD$ = 0.024). However, the proportion of EP relative to the whole cardiac cycle (from one R peak to the next) increased with rising mean heart rate (Pearson's $r(43) = 0.92$, $p < 0.001$), comprising between 0.25 up to 0.45 ($M$ = 0.33, $SD$ = 0.042) of the total mean RR interval (**Fig. S1b right**). This non-proportional shrinking of systolic intervals at higher heart rates underpins the crucial physiological role of systole within the cardiac cycle: Cardiac cycle lengths shorten mainly at cost of diastole to ensure sufficient blood ejection at higher heart rates.

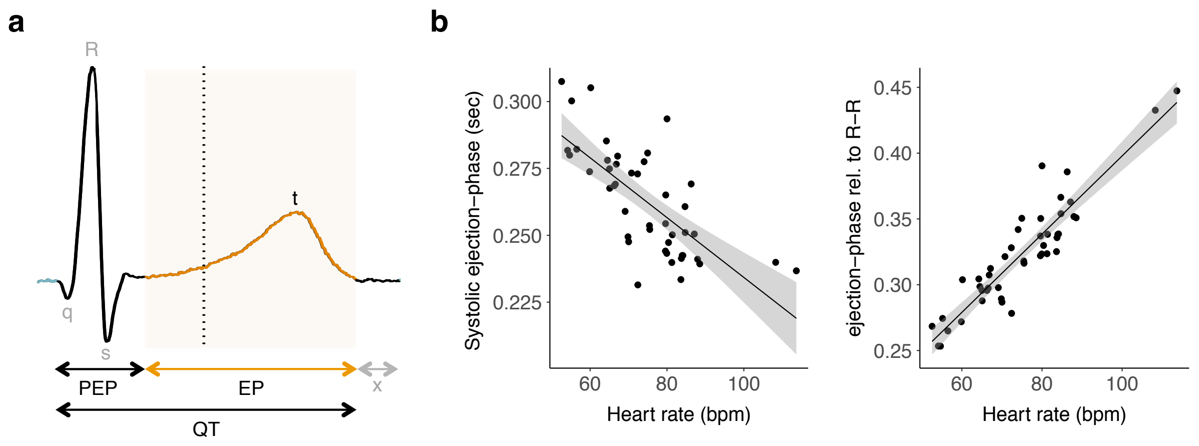


**Figure S1. Extraction of individual ejection phase based on the ECG and inter-individual variation of cardiac phases with heart rate. *a*,** The total systolic interval (onset Q wave until T wave end, QT) was divided into pre-ejection period (PEP) and ejection phase (EP). To prevent an overlap of systole and diastole, a window of 50 ms was inserted after T wave end (indicated by an "x"). Diastole was defined as the remaining part within the RR interval (RR–QT–50 ms). ***b*, left:** Between subjects, the absolute duration of systolic EP decreased with increasing mean heart rate (Pearson's $r$ (43) = -0.72, $p < 0.001$) while **right:** the systolic proportion within the total cardiac cycle (RR interval) increased (Pearson's $r$ (43) = 0.92, $p < 0.001$). That is, cardiac phase lengths shrink disproportionally with faster heartbeats. Grey areas represent the 95% confidence intervals.

2.2 Time ranges of individualized cardiac phases. Participants' systoles (**Fig. S2**) began between 53.0–82.4 ms ($M$ = 65.1, $SD$ = 6.25) after the R peak and ended between 296–375 ms ($M$ = 329, $SD$ = 22.2) after the R peak. Systole duration ranged from 232–308 ms ($M$ = 264, $SD$ = 19.7), also depending on the participant's resting heart rate (cf. **Fig. S1**). These values resemble the time intervals of increased baroreceptor signalling that were reported or used in previous studies: at aortic and carotid sites 90–390 ms after the R peak, with the greatest output around 250 ms (e.g., Edwards, Ring, McIntyre, Winer, & Martin, 2009); at the reticular formation 100 ms after the R peak (Lambertz & Langhorst, 1995); at other brainstem areas 180–270 ms (from the aortic arch) and 210–320 ms (from the carotid sinus) after the onset of ventricular contraction (Dembowsky & Seller, 1995), or 250–350 ms after the R peak (Edwards, McIntyre, Carroll, Ring, & Martin, 2002; Fiacconi, Peter, Owais, &

Köhler, 2016; Gray, Rylander, Harrison, Wallin, & Critchley, 2009). As central processing of baroreceptor feedback signals has often been associated with the T wave (Garfinkel et al., 2013, 2014; Pfeifer et al., 2017) and our systolic time intervals comprise the T wave (i.e., end where the T wave ends), the central representation of cardiac signals can be considered to be included. Participants' mean diastolic phases (**Fig. S2**) started between 346–425 ms (*M* = 379, *SD* = 22.2) after the R peak and ended between 652–1101 ms (*M* = 805, *SD* = 129) after the R peak, thus including earlier used diastolic time points, such as R + 400 ms (Lacey & Lacey, 1974), R + 480 ms (Pramme, Larra, Schächinger, & Frings, 2016), or R + 500 ms (Azevedo, Badoud, & Tsakiris, 2018; Fiacconi et al., 2016; Waselius, Wikgren, Halkola, Penttonen, & Nokia, 2018). Mean diastole duration ranged from 280–678 ms (*M* = 426, *SD* = 113). We conclude that our ECG segmentation allows a physiologically sound approximation of the distinct phases of alternating baroreceptor activity or their central integration – which is comparable with previous studies.

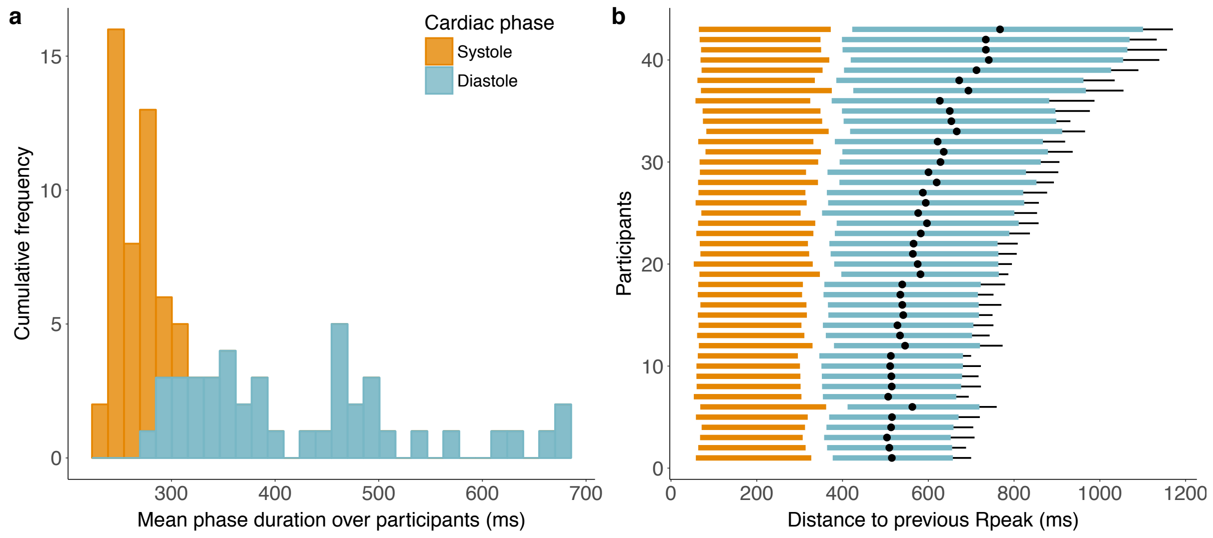


**Figure S2.** Overview of computed cardiac phases ***a,*** the cumulative frequencies of participants' phase lengths for individual systole templates (orange) and mean diastole (blue) together with ***b***, ranges (in relation to the previous R peak) of systole templates (orange) as well as mean diastolic intervals (blue) and their standard deviation (black lines), and average midpoints (black dot) for each participant (ordered by mean diastole length).

3. Supplementary References.


Angell James, J. E. (1971). The effects of altering mean pressure, pulse pressure and pulse frequency on the impulse activity in baroreceptor fibres from the aortic arch and right subclavian artery in the rabbit. *The Journal of Physiology*, *214*(1), 65–88. https://doi.org/10.1113/jphysiol.1971.sp009419

Azevedo, R. T., Badoud, D., & Tsakiris, M. (2018). Afferent cardiac signals modulate attentional engagement to low spatial frequency fearful faces. *Cortex*, *104*(July), 232–240. https://doi.org/10.1016/J.CORTEX.2017.06.016

Boudoulas, H., Geleris, P., Lewis, R. P., & Rittgers, S. E. (1981). Linear relationship between electrical systole, mechanical systole, and heart rate. *Chest*, *80*(5), 613–617. https://doi.org/10.1378/chest.80.5.613

Boudoulas, H., Rittgers, S. E., Lewis, R. P., Leier, C. V., & Weissler, A. M. (1979). Changes in diastolic time with various pharmacologic agents: implication for myocardial perfusion. *Circulation*, *60*(1), 164–169. https://doi.org/10.1161/01.CIR.60.1.164

Coblentz, B., Harvey, R. M., Ferrer, M. I., Cournand, A., & Richards, D. W. (1949). The Relationship Between Electrical and Mechanical Events in the Cardiac Cycle of Man. *British Heart Journal*, *11*(1), 1–22. https://doi.org/10.1136/hrt.11.1.1

Coleridge, H. M., Coleridge, J. C., Poore, E. R., Roberts, A. M., & Schultz, H. D. (1984). Aortic wall properties and baroreceptor behaviour at normal arterial pressure and in acute hypertensive resetting in dogs. *The Journal of Physiology*, *350*(1), 309–326. https://doi.org/10.1113/jphysiol.1984.sp015203

Dembowsky, K., & Seller, H. (1995). Arterial baroreceptor reflexes. In D. Vaitl & R. Schandry (Eds.), *From the Heart to the Brain: The Psychophysiology of Circulation–Brain Interaction.* (pp. 35–60). Europäischer Verlag der Wissenschaften Frankfurt aM.

Edwards, L., McIntyre, D., Carroll, D., Ring, C., & Martin, U. (2002). The human nociceptive flexion reflex threshold is higher during systole than diastole. *Psychophysiology*, *39*(5), 678–681. https://doi.org/10.1017/S0048577202011770

Edwards, L., Ring, C., McIntyre, D., & Carroll, D. (2001). Modulation of the human nociceptive flexion reflex across the cardiac cycle. *Psychophysiology*, *38*(4), 712–718. https://doi.org/10.1017/S0048577201001202

Edwards, L., Ring, C., McIntyre, D., Winer, J. B., & Martin, U. (2009). Sensory detection thresholds are modulated across the cardiac cycle: Evidence that cutaneous sensibility is greatest for systolic stimulation. *Psychophysiology*, *46*(2), 252–256. https://doi.org/10.1111/j.1469-8986.2008.00769.x

Fiacconi, C. M., Peter, E. L., Owais, S., & Köhler, S. (2016). Knowing by heart: Visceral feedback shapes recognition memory judgments. *Journal of Experimental Psychology*, *145*(5), 559–572. https://doi.org/10.1037/xge0000164

Fozzard, H. A. (1977). Heart: excitation-contraction coupling. *Annual Review of Physiology*, *39*(1), 201–220. https://doi.org/10.1146/annurev.ph.39.030177.001221

Fridericia, L. S. (1920). Die Systolendauer im Elektrokardiogramm bei normalen Menschen und bei Herzkranken. *Journal of Internal Medicine*, *53*(1), 469–486. https://doi.org/10.1111/j.0954-6820.1920.tb18266.x

Garfinkel, S. N., Barrett, A. B., Minati, L., Dolan, R. J., Seth, A. K., & Critchley, H. D. (2013). What the heart forgets: Cardiac timing influences memory for words and is modulated by metacognition and interoceptive sensitivity. *Psychophysiology*, *50*(6), 505–512.

https://doi.org/10.1111/psyp.12039

Garfinkel, S. N., Minati, L., Gray, M. A., Seth, A. K., Dolan, R. J., & Critchley, H. D. (2014). Fear from the Heart: Sensitivity to Fear Stimuli Depends on Individual Heartbeats. *Journal of Neuroscience*, *34*(19), 6573–6582. https://doi.org/10.1523/JNEUROSCI.3507-13.2014

Gill, H., & Hoffmann, A. (2010). The timing of onset of mechanical systole and diastole in reference to the QRS-T complex: A study to determine performance criteria for a non-invasive diastolic timed vibration massage system in treatment of potentially unstable cardiac disorders. *Cardiovascular Engineering*, *10*(4), 235–245. https://doi.org/10.1007/s10558-010-9108-x

Gray, M. A., Rylander, K., Harrison, N. A., Wallin, B. G., & Critchley, H. D. (2009). Following One's Heart: Cardiac Rhythms Gate Central Initiation of Sympathetic Reflexes. *Journal of Neuroscience*, *29*(6), 1817–1825. https://doi.org/10.1523/JNEUROSCI.3363-08.2009

Lacey, B. C., & Lacey, J. I. (1974). Studies of heart rate and other bodily processes in sensorimotor behavior. In P. A. Obrist, A. H. Black, J. Brener, & L. V. DiCara (Eds.), *Cardiovascular psychophysiology: Current issues in response mechanisms, biofeedback and methodology.* (pp. 538–564). New Brunswick, NJ, US: AldineTransaction. Retrieved from http://psycnet.apa.org/record/2007-11499-026

Lambertz, M., & Langhorst, P. (1995). Cardiac rhythmic patterns in neuronal activity are related to the firing rate of the neurons: I. Brainstem reticular neurons of dogs. *Journal of the Autonomic Nervous System*, *51*(2), 165–173. https://doi.org/10.1016/0165-1838(94)00128-7

Levick, J. R. (2010). *An introduction to cardiovascular physiology* (5th ed.). London: Hodder Education Publishers.

Lewis, R. P., Rittogers, S. E., Froester, W. F., & Boudoulas, H. (1977). A critical review of the systolic time intervals. *Circulation*, *56*(2), 146–158. https://doi.org/10.1161/01.CIR.56.2.146

Lewis, T. (1920). The Mechanism and Graphic Registration of the Heart Beat. *JAMA: The Journal of the American Medical Association*, *75*(15), 1019. https://doi.org/doi: 10.1001/jama.1920.02620410049029

Lombard, W. P., & Cope, O. M. (1926). The duration of the systole of the left ventricle of man. *American Journal of Physiology--Legacy Content*, *77*(2), 263–295. Retrieved from https://doi.org/10.1152/ajplegacy.1926.77.2.263

Pappano, A. J., & Wier, W. G. (2013). *Cardiovascular physiology* (10th ed.). Elsevier/Mosby. Retrieved from https://www.sciencedirect.com/book/9780323086974/cardiovascular-physiology

Pfeifer, G., Garfinkel, S. N., Gould van Praag, C. D., Sahota, K., Betka, S., & Critchley, H. D. (2017). Feedback from the heart: Emotional learning and memory is controlled by cardiac cycle, interoceptive accuracy and personality. *Biological Psychology*, *126*, 19–29. https://doi.org/10.1016/j.biopsycho.2017.04.001

Pramme, L., Larra, M. F., Schächinger, H., & Frings, C. (2016). Cardiac cycle time effects on selection efficiency in vision. *Psychophysiology*, *53*(11), 1702–1711. https://doi.org/10.1111/psyp.12728

Quelhas Martins, A., McIntyre, D., & Ring, C. (2014). Effects of baroreceptor stimulation on performance of the Sternberg short-term memory task: A cardiac cycle time study. *Biological Psychology*, *103*, 262–266. https://doi.org/10.1016/j.biopsycho.2014.10.001

Rushmer, R. F. (1976). *Cardiovascular dynamics* (4th ed.). London: W.B. Saunders.

Sherwood, A., Allen, M. T., Fahrenberg, J., Kelsey, R. M., Lovallo, W. R., & van Doornen, L. J. P. (1990). Methodological Guidelines for Impedance Cardiography. *Psychophysiology*, *27*(1), 1–23. https://doi.org/10.1111/j.1469-8986.1990.tb02171.x

Vázquez-Seisdedos, C. R., Neto, J. E., Marañón Reyes, E. J., Klautau, A., de Oliveira, R. C., & Limão de Oliveira, R. C. (2011). New approach for T-wave end detection on electrocardiogram: Performance in noisy conditions. *BioMedical Engineering OnLine*, *10*(1), 77. https://doi.org/10.1186/1475-925X-10-77

Wallace, A. G., Mitchell, J. H., Skinner, N. S., & Sarnoff, S. J. (1963). Duration of the Phases of Left Ventricular Systole. *Circulation Research*, *12*(6), 611–619. https://doi.org/10.1161/01.RES.12.6.611

Waselius, T., Wikgren, J., Halkola, H., Penttonen, M., & Nokia, M. S. (2018). Learning by heart: cardiac cycle reveals an effective time window for learning. *Journal of Neurophysiology*, *120*(2), 830–838. https://doi.org/10.1152/jn.00128.2018

Weissler, A. M., Harris, W. S., & Schoenfeld, C. D. (1969). Bedside technics for the evaluation of ventricular function in man. *American Journal of Cardiology*, *23*(4), 577–583. https://doi.org/10.1016/0002-9149(69)90012-5

Weissler, A. M., Harris, W. S., & Schoenfield, C. D. (1968). Systolic Time Intervals in Heart Failure in Man. *Circulation*, *37*(2), 149–159. Retrieved from http://circ.ahajournals.org/cgi/content/abstract/37/2/149

Wiggers, C. J. (1921). Studies on the Consecutive Phases of the Cardiac Cycle: I. The Duration of the Consecutive Phases of the Cardiac Cycle and the Criteria for Their Precise Determination. *American Journal of Physiology--Legacy Content*, *56*(3), 415–438. https://doi.org/10.220.33.3

Wiggers, C. J. (1923). Modern Aspects of the Circulation in Health and Disease. *The Journal of the American Medical Association*, *81*(15), 1305. https://doi.org/10.1001/jama.1923.02650150059033
